# Development of cell-free transcription-translation systems in three soil Pseudomonads

**DOI:** 10.1101/2023.06.09.544292

**Authors:** Joseph T. Meyerowitz, Elin M. Larsson, Richard M. Murray

## Abstract

*In vitro* transcription-translation (TX-TL) can enable faster engineering of biological systems. This speed-up can be significant, especially in difficult-to-transform chassis. This work shows the successful development of TX-TL systems using three soil-derived wild-type Pseudomonads known to promote plant growth: *Pseudomonas synxantha, Pseudomonas chlororaphis*, and *Pseudomonas aureofaciens*. One, *P. synxantha*, was further characterized. A lysate test of *P. synxantha* showed a maximum protein yield of 2.5 *μ*M at 125 proteins per DNA template and a maximum protein synthesis rate of 20 nM/min. A set of different constitutive promoters driving mNeonGreen expression were tested in TX-TL and integrated into the genome, showing similar normalized strengths for *in vivo* and *in vitro* fluorescence. This correspondence between the TX-TL derived promoter strength and the *in vivo* promoter strength indicates these lysate-based cell-free systems can be used to characterize and engineer biological parts without genome integration, enabling a faster designbuild-test cycle.

## Introduction

The field of synthetic biology has advanced considerably over the past years. From scar-free DNA assembly [1] to CRISPR-mediated genome editing [2], new technologies have made it possible to tackle increasingly complex tasks in the engineering and control of biological systems. Synthetic biology has advanced to a point where we can reliably engineer bacteria to perform varying and complex tasks including synthesis of precious chemicals [3, 4], performance of logical operations [5, 6] and sensing [7].

Substantial challenges still remain with the use of synthetic biology tools. While the cost of DNA sequencing and DNA synthesis continues to decline [8, 9], and databases of genetic information [10], protein structure [11], and characterized genetic parts [12, 13, 14] continue to expand, composing these separate advances into engineered biological systems is complicated and incompletely systematized. Because of this difficulty, new techniques in the field are often implemented only in well-understood model organisms like *E. coli, B. subtilis, P. putida* and *S. cerevisiae*. To expand the range of projects possible with synthetic biology, developing tools for use with non-model organisms is essential. Environmental bacteria often have desirable traits unavailable in model organisms, such as the ability to perform specific types of metabolism [15], tolerate certain stresses and inhibitors [16] and colonize different environments [17, 18, 19]. Making non-model organisms tractable for engineering also enables targeted experiments to reveal the roles these organisms play in their natural niches.

While some advances have been made in engineering non-model organisms [20, 21, 22], “domesticating” a non-model organism remains challenging. For a new organism of interest, the first steps for engineering are to find methods for cultivation and transformation. A later step is to characterize genetic parts from which new circuits can be built. One reason why working with non-model organisms is challenging comes from the variation in each species’ underlying biology, requiring species-by-species tailoring of every protocol. An example of this variation can be seen in the mechanism for ribosomal initiation, which can have dramatic differences from commonly used organisms such as *E. coli* which use Shine-Delgarno led initiation and other bacteria such as *Deinococcus deserti* or *Mycobacterium tuberculosis* which frequently use leaderless mRNA sequences without a Shine-Delgarno sequence [23].

One approach used to accelerate engineering of non-model organisms is *in vitro* transcription-translation extract (TX-TL), or “cell-free”. TX-TL reactions re-comprise the machinery necessary for RNA transcription and protein translation. By removing soluble internal components from a species of interest and producing a clarified lysate, this process results in a non-living, simplified system that can be used to prototype genetic parts and enzymes. TX-TL is particularly useful for the study of non-model organisms that may be difficult to genetically transform or grow under certain conditions. Instead of cloning and re-isolating the correct transformed cells, plasmid DNA or PCR products containing the circuit can be added to a mixture of the lysate and other components needed such as nucleotides and salts. This procedure allows characterization of the test parts within hours. Genetic manipulation of some non-model organisms can take days or weeks to complete.

Creating TX-TL reactions using extracts from non-model organisms can be challenging, as the activity depends on factors such as growth phase during harvest. Some species have long doubling times which makes the process of reaching high cell densities time consuming and contamination-prone. Because of the extensive knowledge about the *E. coli* genome, certain genetic alterations have been shown to increase *in vitro* activity by removing DNA and amino acid degrading enzymes [24]. Such a strategy cannot be applied generally with non-model organisms, as the identify of similar genes may not be known. Despite these challenges, TX-TL has been successfully created from non-model organisms including *B. megaterium* [25] and several Streptomycetes [26] .

In this study, we demonstrate the production and characterization of TX-TL from three soil-derived wild-type Pseudomonads. This class of organisms is important because of their ecological ubiquity. These species are known for promoting plant growth by releasing phenazines, an antimicrobial class of molecules, into the environment [27, 28]. These phenazines help protect food crops from disease. It has also been suggested that these molecules may play a role in nutrient cycling in the soil [29]. Pseudomonads also demonstrate an innate tolerance to certain growth inhibitory compounds in the environment and in industrial fermentation settings. Their ability to tolerate solvents and certain growth inhibitors found in low-cost industrial feedstock make them good candidates for commercial applications [30]. These bacteria can sometimes metabolize lignocellulosic growth inhibitors as well, showcasing their evolutionary adaptation to environments abundant in plant material [31, 32]. Root-living bacteria typically also evolve tolerance to antimicrobial compounds, such as penicillins, produced by other bacteria competing to inhabit the same ecological niche [33]. The creation of these three extracts from wild-type strains expand the toolkit for engineering these and other environmental bacteria.

## Results and Discussion

Soil provides a high-lignin environment where soil bacteria may have evolved tolerance to certain stresses ill-tolerated by *E. coli* [34]. To find potential new chassis for engineering with the desired stress tolerances, three soil-derived bacteria, *Pseudomonas synxantha* 2-79 (“PSX”), *Pseudomonas chlororaphis* PCL1391 (“PCL”), and *Pseudomonas aureofaciens* 30-84 (“PAU”), were chosen for testing.

These *Pseudomonas* species have been studied separately and together as symbionts of plant roots, where they produce anti-fungal phenazines [35, 36, 37]. The phenazines from these bacteria are known to control take-all, a disease in wheat plant roots, and damping-off disease, a multi-pathogen constellation of seed and seedling injuries across different plant species [38]. *P. aureofaciens* 30-84 is also called *P. chlororaphis subvar. aureofaciens* 30-84, and is closely related to *P. chlororaphis* PCL1391. PSX is more distantly related, with a similar distance between its genome, *E. coli*, PCL, and PAU as measured by digital DNA-DNA hybridization [39]. The phylogenetic relationships between the three target bacteria and other bacteria used for TX-TL are described in Supplementary Information A. All three of the Pseudomonads are wild-type isolates without many of the common mutations such as restriction system knockouts seen in more domesticated strains [40].

### Pseudomonads show phenotypic differences from the *E. coli* BL21 Rosetta2 used in traditional TX-TL

To confirm the expected tolerance of some environmentally and industrially relevant compounds, a series of *in vivo* growth experiments were done to compare the growth ability of the Pseudomonads and a reference *E. coli* BL21 Rosetta2 strain often used in *E. coli* TX-TL. First, the three Pseudomonads were grown in the presence of the lignocellulosic monomer syringaldehyde. PSX showed tolerance at all tested concentrations, wheras PCL and PAU had minimal difference in their tolerance compared to *E. coli* (Figure 1A). Next, we compared the tolerance of three common antibiotics (two ribsomal inhibitors and one cell-wall synthesis inhibitor) for PSX and *E. coli* (Figure 1B-D). As expected PSX, which has evolved to tolerate compounds frequently present in the rhizosphere, show higher tolerance than *E. coli*.

**Figure 1:**
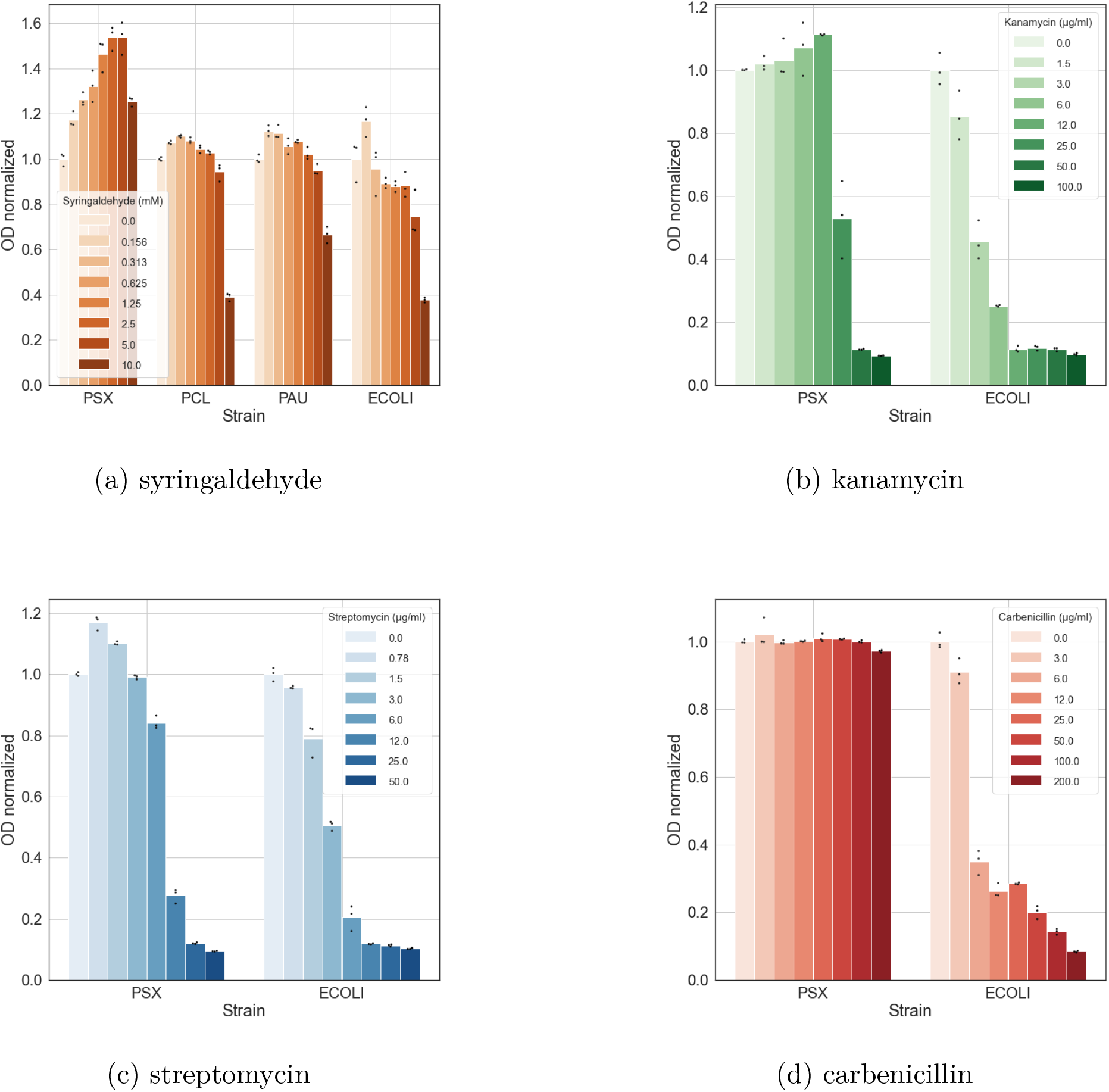
Differences in growth when exposed to (A) syringaldehyde, (B) kanamycin, (C) streptomycin and (D) carbenicillin. Barplots are showing one biological replicate of the final optical density (OD) after 24 hours of growth normalized to the OD of cells grown without any growth inhibitor.

These results confirm that at least one of the tested Pseudomonads has meaningful phenotypic differences from the standard *E. coli* for engineering purposes. The ability of PSX to broadly tolerate different types of stress is promising for later uses in bioproduction or as a biocontrol agent for agriculture.

### *In vitro* transcription-translation systems are possible with three new wild-type Pseudomonads

After the tolerance experiments, we continued towards the effort of making cell-free extracts from the Pseudomonads with the aim of achieving functional extracts that can be compared to *E. coli* extract in terms of productivity.

We started scale-up by growing 50 mL cultures in 250 mL baffled flasks of each species with inoculum sizes of 1:100 and 1:1000 from overnight 5 mL cultures in 2xYTPG. Growth was sufficient to support production of cell pellets for TX-TL use, shown in Supplementary Information B, and identified likely time and OD600 ranges for early exponential phase harvest.

Scale-up continued to 660 mL scale in 2.8L baffled flasks. Cultures were harvested at OD 3.0 for all three *Pseudomonas* species to begin with; initial testing showed *P. chlororaphis* lysate required an earlier harvest, chosen to be at OD 1.5 (data not shown). Each replicate for each species was grown on a different day in 3x 660 mL volume, harvested, washed, and frozen in two 50 mL tubes.

Lysates were produced using one of the two frozen pellets with varying process conditions. In previous work, two of the most important parameters to optimize after harvest time have been lysis and runoff [41]. Here we try three different sonication amplitudes and three different runoff times, and test each combination across six different reaction conditions (Figure 2A). These varied protocol parameters show different protein yields across lysates from all three species (Figure 2B).

**Figure 2:**
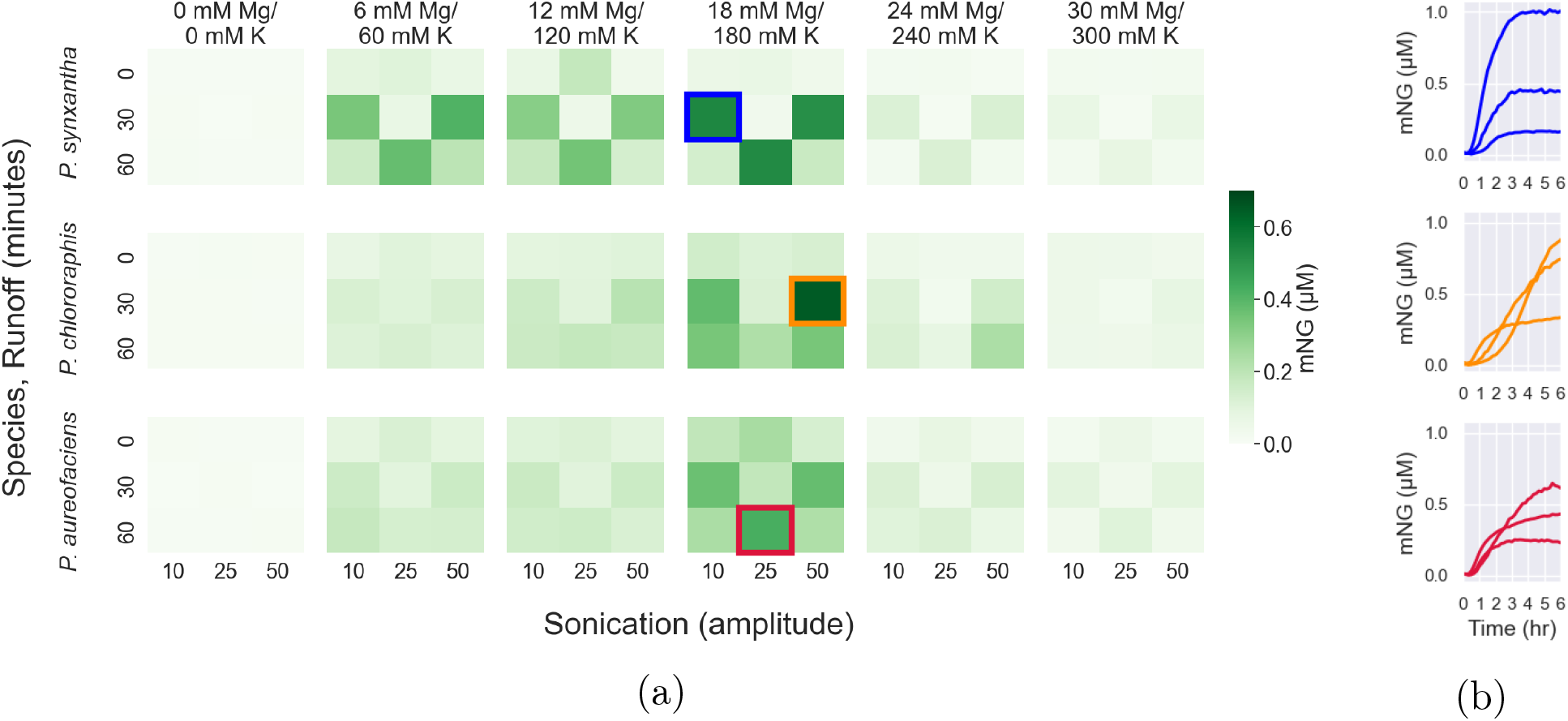
All three species show productive TX-TL systems across a range of lysate preparation and reaction parameters. (a) Mean protein yield across biological replicates, different sonication amplitudes, runoff times, salt concentrations, and species, shown after 6 hours of TX-TL reaction time. (b) Individual time traces from each species showing biological replicates.

One important note is that these reaction conditions were prepared by creating a single mix of energy solution, amino acid mix, and PEG, then adding it to 384 well plates with the different salts. The pattern of increased and decreased yield is unlikely to be the result of differences in the preparation of the reaction buffer, as all of the data here come from the preparation of a single mixture of the common reaction components.

### Repeating lysis production at larger scale improved yield and consistency of *P. synxantha* TX-TL

Next, three new “big batches” of PSX lysate were produced on separate days at a single sonication amplitude (‘50’) and a single runoff time (60 minutes). These conditions were chosen based on results from the first set of replicates from the sonication and runoff testing, and were not at the overall maximum yield shown across the biological triplicates above. These “big batches” used the second of the two frozen pellets from the triplicate growths described above. Batch 1 was made from the same harvested culture as the first PSX replicate in Figure 2, Batch 2 and Batch 3 correspond to the second and third cultures used as well.

These “big batches” show an approximately doubled protein yield and increased consistency between biological replicates compared to the reactions in Figure 2 and were produced from the same cultures. There were improvements in speed of processing time due to the simpler process conditions and increased experience with processing the pellets into lysates. The causes of the improvements in these follow-on lysates were not explored in greater depth.

Potassium and magnesium salts were tested with more detail across a narrower range of concentrations (Figure 3A). The test conditions show a clear fluorescent signal accumulating over the first 4-5 hours of the reaction for all three of the new batches (Figure 3B). The yields and optimum salt concentrations vary across the different batches to a degree similar to past *E. coli* TX-TL reactions. These results demonstrate the extract making process was repeatable, and that there are some differences in performance between the batches despite being made with the same process parameters.

**Figure 3:**
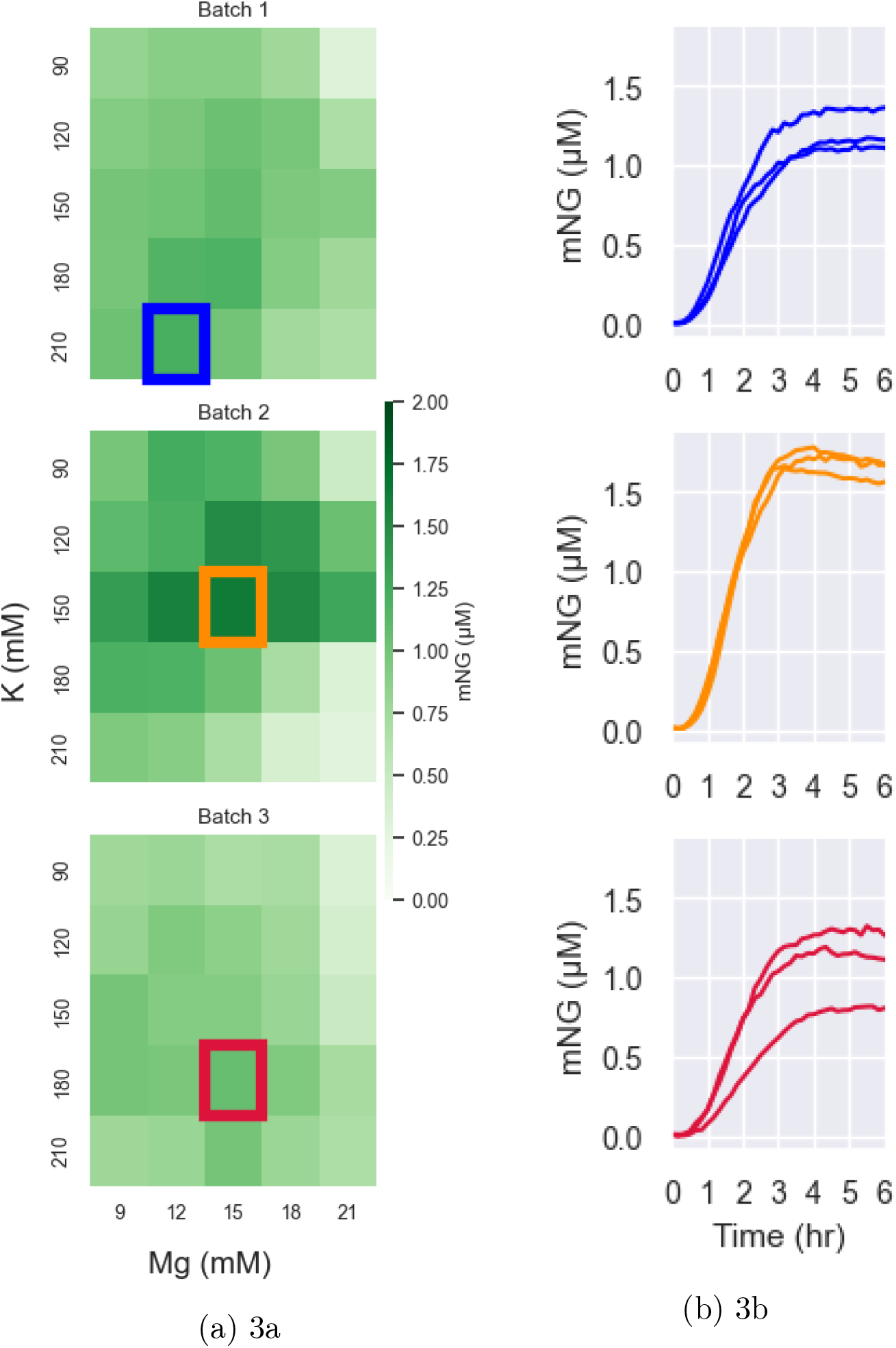
Salt panel for the big batches of PSX extract. (a) Variations in reaction salt optima across three batches of *P. synxantha* lysate shown after 6 hours of TX-TL reaction time. (b) Individual time traces from each batch, each chart showing technical replicates.

### Increases in DNA template produce higher protein yields

Next, using these three “big batches” of extract with their respective salt optimization values, we tested a varying amount of DNA template added to the reaction (Figure 4A-B). Each batch of extract produced a similar amount of the fluorescent reporter for a given template concentration. Across the tested template concentrations, more template always results in more protein. Doubling the amount of template doubles the amount of fluorescent reporter made within some of the tested range.

**Figure 4:**
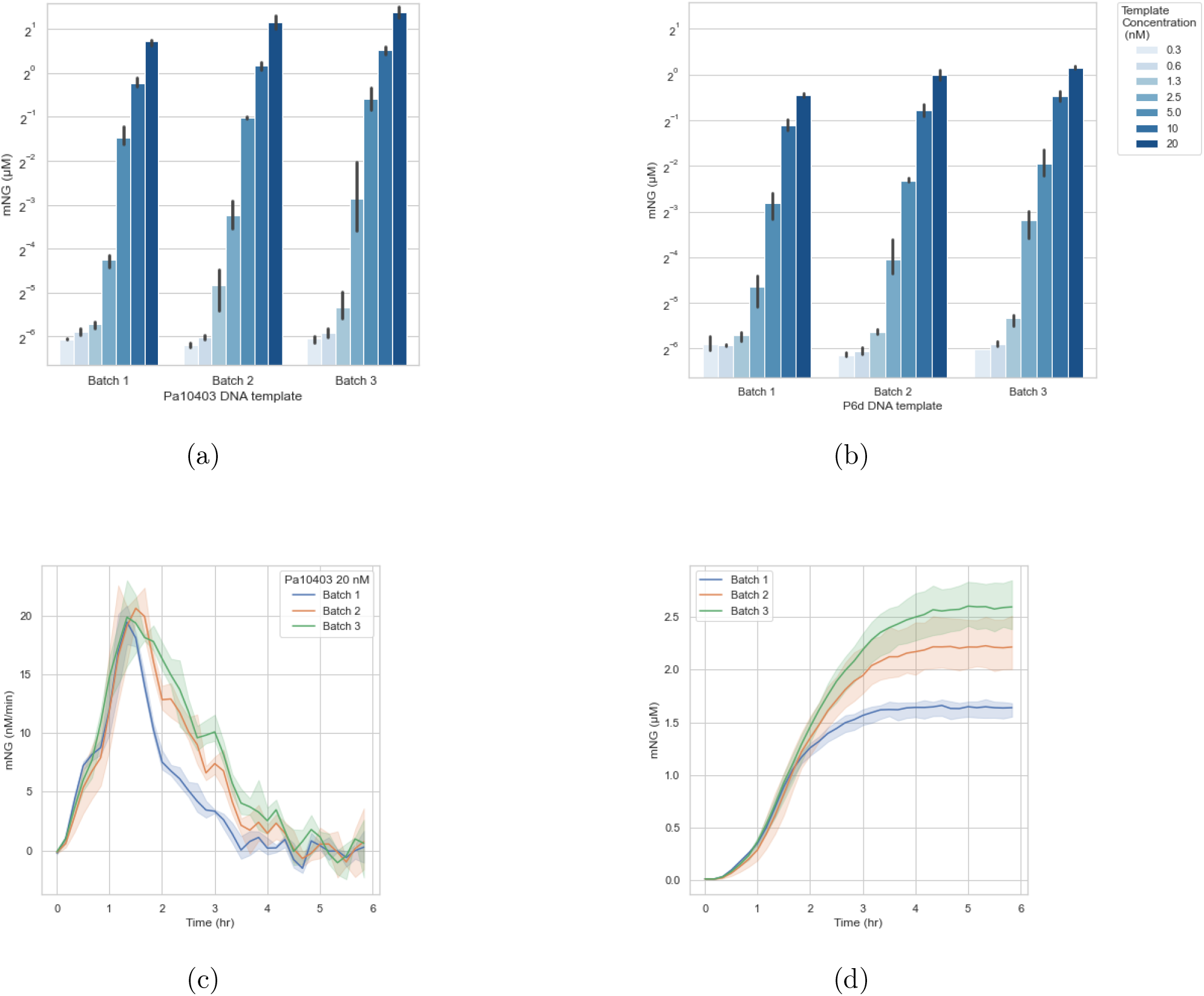
Fluorescence measurements at different DNA template concentrations. (a and b) Endpoint protein yield across the three *P. synxantha* extract batches with varied DNA template concentration and two different templates. (c and d) Protein synthesis rates and cumulative protein yield across the same three batches with Pa10403 template at 20.0 nM. Error bands show standard deviation across technical replicates.

The 20 nM Pa10403 condition using the third extract batch reached the highest concentration of mNeongreen among all of the PSX experiments here at approximately 2.5 *μ*M with a peak synthesis rate of 20 nM/min (Figure 4C-D). Over the course of the reaction this protein yield is equal to 125 proteins produced per DNA template.

Qualitatively all of the extracts stop producing mNeonGreen at around 4 hours regardless of the amount of template that is added. In some circumstances we might expect more heavily burdened TX-TL reactions to exhaust some key resource earlier than less burdened reactions, but that is not what we see here. Protein synthesis rates are similar across all three batches in the early part of the reaction through the peak. The differences in protein yield between the extracts are attributable to different rates of the reaction slowing post-peak.

### Promoters tested *in vivo* and *in vitro* show similar strengths

To evaluate whether we could use the *P. synxantha* extract for prototyping of genetic elements as seen in previous studies of *E. coli* [42] and *B. megaterium* [25], we compared protein synthesis levels for 11 different constitutive promoters from *E. coli* and *P. putida* driving expression of a genomically integrated fusion with the green fluorescent protein mNeonGreen (Figure 5).

**Figure 5:**
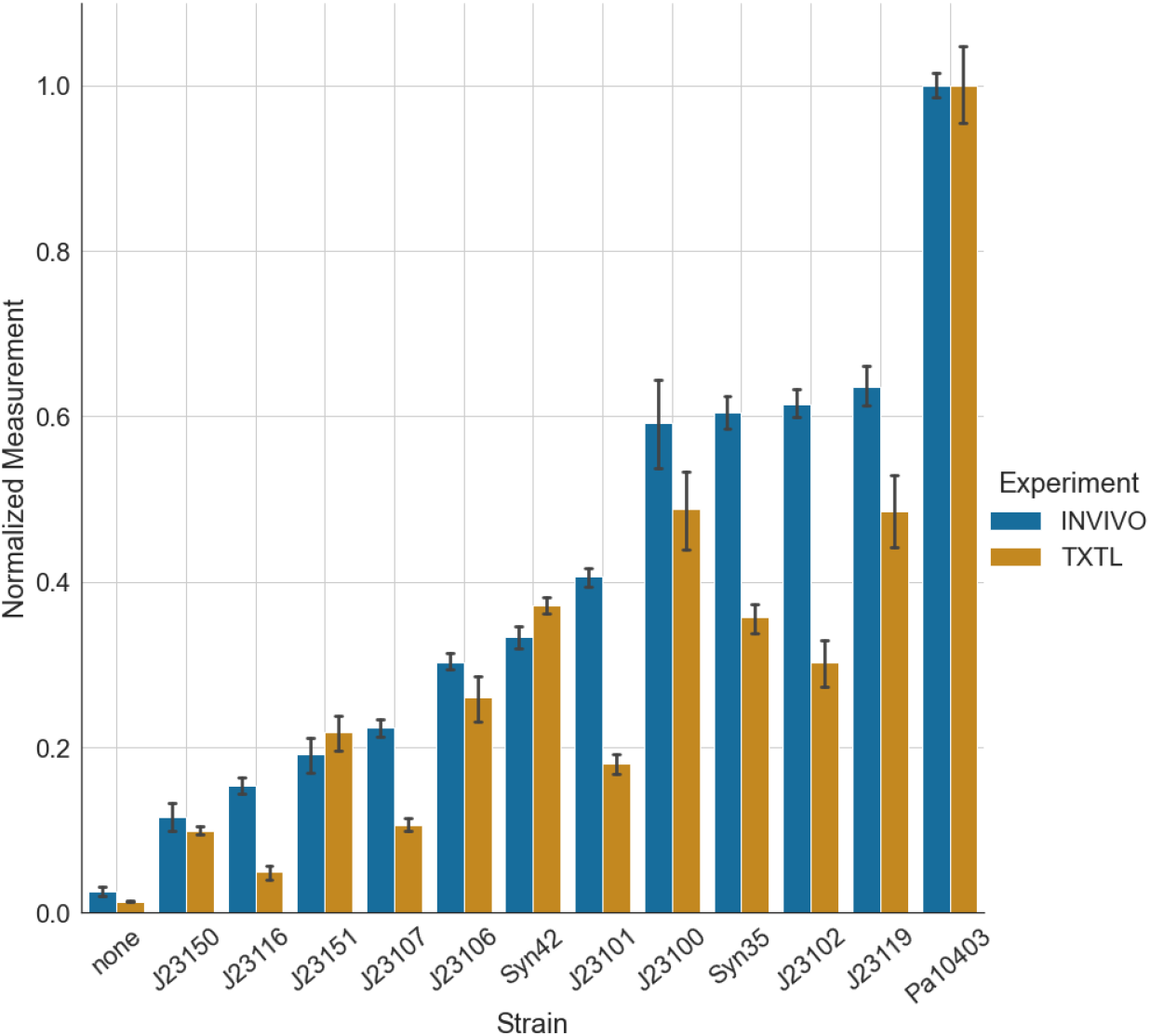
Constitutive promoters driving the expression of mNeonGreen were compared *in vivo* in *P. synxantha* living cells and *in vitro* in *P. synxantha* cell-free extract. All expression levels are normalized to the expression level of the strongest promoter fusion using the Pa10403 promoter.

The fluorescence below is normalized by optical density at 500 nm at 24 hours of growth. To compare the results from the *in vitro* TX-TL reactions, we take the fluorescence from a time course at 6 hours into the reaction. We then normalize all fluorescence values to the largest fluorescence value of the strongest promoter Pa10403 [43]. The reactions *in vitro*, regardless of promoter strength, all stop producing protein at approximately the same time.

We can distinguish between strong and weak constitutive promoters using cell-free prototyping, but we cannot always accurately predict the rank of strength within these groups, as seen in previous work [25]. The normalized fluorescence in the TX-TL reactions is consistently lower than the corresponding *in vivo* values. Even though the *in vivo* to TX-TL comparison is not perfectly matched, the number of genetic elements that need to be tested *in vivo* can be reduced by first prototyping them using the TX-TL system. This also highlights the limitation of using TX-TL as a prototyping tool for *in vivo* expression.

## Conclusion

Together these experiments show that our target *Pseudomonas* spp. have pre-existing tolerance to some growth inhibitors that reduce the growth capacity of *E. coli*, and these species are tractable for further engineering work. The TX-TL measurements of the promoter panel match the *in vivo* measurements of the same promoters. This shows the TX-TL system can be used for characterizing parts for later use *in vivo*. The ability to perform TX-TL reactions with these species is a key component of engineering and rapid prototyping with new non-model organisms. A working TX-TL system can enable a faster design-build-test cycle, and make characterization of new parts practical.

## Methods

### Bacterial strains and growth conditions

*Pseudomonas synxantha* 2-79, *Pseudomonas chlororaphis* PCL1391, and *Pseudomonas aureofaciens* 30-84 were obtained from the Newman lab at Caltech. Cells were grown by streaking onto 2xYTPG (16 g/L tryptone, 10 g/L yeast extract, 5 g/L NaCl, 40 mM potassium phosphate dibasic, 22 mM potassium phosphate monobasic, 2% glucose) agar or LB (10 g/L tryptone, 5 g/L yeast extract, 5 g/L NaCl) agar without antibiotics and incubating for 18-36 hours at 30°C. Individual colonies were picked and grown in 5 mL of 2xYTPG with shaking at 200-220 rpm at 30°C overnight. As needed, 50 mL of 2xYTPG in a 250 mL baffled flask or 660 mL of 2xYTPG in a 2.8L baffled flask with an adhesive AirOTop sterile seal would be inoculated 1:1000 the next day. Cell growth was monitored by optical density at 500 nm or 600 nm, with 1:1, 1:4, 1:16, 1:64, and 1:256 dilutions of culture samples prepared in fresh 2xYTPG media to stay in the Biotek H1MF plate reader linear range for absorption measurements (OD 0.005 to 0.5).

### *In vivo* growth inhibition tests

Cells were grown with continuous shaking in a Biotek H1MF plate reader for 24 hours. The cells were grown in LB medium with added growth inhibitor in serial dilutions with a starting concentration of 10 mM syringaldehyde, 50 *μ*g/ml streptomycin, 100 *μ*g/ml kanamycin and 200 *μ*g/ml carbenicillin. *P. synxantha* was grown at 30°C and *E. coli* was grown at 37°C.

### Construction of DNA templates

DNA parts including promoters, ribosome binding sites, reporters, terminators, and backbones with antibiotic resistance were amplified by PCR in NEB Q5 2x Master Mix, purified by gel extraction and QIAGEN MinElute spin columns, and assembled using NEB Hifi 2x Master Mix. Supplemental Information F shows the plasmid map for a strong constitutive promoter expressing mNeonGreen, and the plasmid map for a promoter with a different strength also expressing mNeonGreen.

### Measurement of protein concentration

For each extract produced for the sonication, runoff, and salt panels, protein concentration was measured using a Bradford assay. In brief, a 1:50 dilution of lysate was prepared in Tris-buffered saline. A bovine serum albumin standard was used to prepare 8 different concentrations for a standard curve. Standard curves were prepared in triplicate on every 96-well plate used for protein measurement, and samples were also prepared in triplicate.

### Production of *Pseudomonas* cell-free extracts

The protocol from Sun et al [36] and the protocol from Kwon et al [37] for *E. coli* TX-TL was repurposed for the *Pseudomonas* TX-TL protocol with minor modifications.

An illustrated and expanded version of our protocol is shown in Supplement X. Briefly, cells are grown on 2xYTPG agar. After 18-36 hours of growth at 30°C, the plates are stored at 4C for up to a month. Individual colonies are picked and used to grow 5 mL overnight cultures in 14 mL tubes. 2.8L baffled shake flasks with 660 mL of 2xYTPG are inoculated 1:1000 the next day and grown for 8-12 hours. Note that unlike past protocols, the final culture is inoculated directly from a 5 mL overnight, without an intermediate 50 mL culture, with a smaller inoculum.

The larger culture is grown to the harvest optical density (OD 1.5 +/-10% for *P. chlororaphis*, OD 3.0 +/-10% for *P. synxantha* and *P. aureofaciens*). Optical density is measured once per 60 minutes until the last doubling of the growth, then monitoring increases to once per 20 minutes. Dilutions of 1:4, 1:16, 1:64, or 1:256 in fresh 2xYTPG media are used to produce a sample with a 600nm absorption over 0.05 and under 0.5. 200 mL of Buffer A (1.8 g/L Tris-acetate, 3 g/L Mg-acetate, and 12.2 g/L K-glutamate, adjusted to pH 8.2 with 2M Tris and autoclaved, then 1 mL / L of 1M DTT is added) is frozen in 1L centrifuge tubes stored at -20C at an angle for 1-2 hours.

At harvest time, the cell culture is decanted into the 1L centrifuge tube over the ice. This brings the culture temperature down from 30°C to 10°C within a few minutes. The cell culture is centrifuged at 4800g for 12 minutes at 4°C. The supernatant is decanted, and the tubes are placed on ice. The cell pellet is resuspended in Buffer A, then otherwise follows the Sun et al. *E. coli* protocol with two wash steps in 1L bottles, a final wash step and transfer into two weighed 50 mL tubes, flash freezing in liquid nitrogen, and stored at -80°C.

At a later time, the cell pellet is removed from the freezer and placed on wet ice to thaw. 1 mL / g of Buffer A is used to resuspend the pellet, then 4 mL of resuspended cell pellet is added to each 14 mL falcon tube. Each tube is sonicated on wet ice for 120s, 5s on, 10s off, with an amplitude of 10, 25, or 50 on a Qsonica Q700 with a 1/8” stepped microtip and microtip coupler. The tubes are centrifuged at 12,000g for 10 minutes at 4°C. The supernatant is transferred to new 1.4 mL or 14 mL tubes and incubated in a “runoff” step with open lids shaking at 220 rpm at 30C for 0-60 minutes. These tubes are centrifuged again at 12,000g for 10 minutes at 4°C, then the supernatant is transferred to new tubes for freezing in LN2 and storage at -80C.

### TX-TL reaction conditions

Reaction conditions also follow the *E. coli* protocol in Sun et. al. (34). In brief, each 10 microliter reaction in a 384 well plate contains 29% buffer, 13% potassium and magnesium glutamate salts, 25% DNA template, and 33% processed lysate. The buffer consists of 26% energy solution, 57% amino acid mix, and 17% 40% PEG-8000. These are mixed on ice and hand-pipetted carefully to avoid introducing bubbles. Individual reactions are prepared in glass-bottomed 384 well plates sealed with an oxygen-permeable Breathe-Easy seal. The plate is centrifuged for 2 minutes to mix the reagents and reduce bubbles in the wells, then read at 30°C in the plate reader.

### Preparation of DNA template for TX-TL reactions

*E. coli* JM109 or DH10B cells with DNA constructs were grown to 100 mL (midiprep) or 400 mL (maxiprep) scale. Pellets were processed using Macherey-Nagel midiprep or maxiprep kits to produce 100s to 1000s of micrograms of purified DNA, including a “finalizer” step to re-purify and concentrate the eluate from the kit. DNA is diluted to produce a concentration of 10 nM in the final reaction unless otherwise specified.

### Measurement of *in vitro* fluorescence

Biotek H1M and H1MF plate readers were used to measure fluorescence of mNeon-Green (excitation 490 nm, emission 520 nm) and optical density (500 nm and 600 nm). mNeonGreen protein was purified using a his-tag and NiNTA columns, and UV absorption was used to quantify the amount of purified protein. Triplicate dilutions at 8 concentrations from 20 nM to 0 nM in 1xPBS were measured five times in all plate readers, and separate standard curves were created for each plate reader. Dilutions were incubated at 30°C during fluorescence measurements.

### Construction of constitutively fluorescent *P. synxantha*

All cloning to produce *P. synxantha* genomic integration constructs was done using

*E. coli* DH10B with the backbone pJM220 [44]. Constitutive promoters from *E. coli* (http://parts.igem.org/Promoters/Catalog/Anderson) and *P. putida* [45] were fused to the fluorescent protein mNeonGreen. The constructs were then integrated on the *P. synxantha* chromosome using transposase based insertion at the Tn7 site. The protocol used for making and transforming competent cells was modified from Choi et al [46].

Briefly, electrocompetent *P. synxantha* cells were electroporated in 1 mm-gap cuvettes (at 1.8 mV, 600 Ω and 10 *μ*F) with the construct plasmid as well as a plasmid containing the transposase and genes required for genome insertion [47]. The cells were then recovered in rich medium (SOC) for 3 hours at 30°C and plated onto LB agar plates containing gentamicin (20 *μ*g/ml) and incubated for 24 hours before picking colonies for sequence verification.

### Plate reader assay for *in vivo* fluorescence

*In vivo* fluorescence was measured using a Biotek (Synergy H1) plate reader. The experiments ran for 24 hours at 30°C using continuous orbital shaking starting from an overnight culture diluted to approximately OD 0.1 into LB medium. OD was measured every 10 minutes at 500 nm and fluorescence was measured at 490/520 nm.

## Author Information

Current address is 1200 E. California Blvd, MC 138-78, Pasadena, CA 91125. The authors declare no competing interests.

## Author Contribution

JTM, EML and RMM conceptualized the project. JTM and EML designed the experiments and analyzed the data. JTM performed TX-TL experiments. EML performed *in vivo* experiments, with the exception of the syringaldehyde experiment which was performed by JTM. EML did the plasmid construction and integration. JTM and EML wrote the manuscript with input from RMM.

## Acknowledgements

With thanks to M. Prator for performing the Bradford assays of the lysates, to Z. Jurado for purifying the mNeonGreen fluorescence standard, to Prof. Dianne Newman for the *Pseudomonas* strains and R. Alcalde for the Pa10403-mNeonGreen plasmid and *P. synxantha* strain, to Dr. R. Sidney Cox III for assistance with data visualization, to A. Pandey for assistance with data processing, and to Dr. Dmitri Mavrodi for assistance with the genomic integration protocol for *P. synxantha*. This research is supported by the Institute for Collaborative Biotechnologies through contract W911NF-19-D-0001, cooperative agreement W911NF-19-2-0026, and grant W911NF-09-0001 from the U.S. Army Research Office, the National Science Foundation through grant CBET-1903477, and the International Human Frontiers Science Program. The content of this paper does not necessarily reflect the position or the policy of the U.S. Government, and no official endorsement should be inferred.

## References

[1] D. G. Gibson, L. Young, R.-Y. Chuang, J. C. Venter, C. A. Hutchison III, and H. O. Smith, “Enzymatic assembly of DNA molecules up to several hundred kilobases,” Nature methods, vol. 6, no. 5, pp. 343–345, 2009.

[2] R. Barrangou and J. A. Doudna, “Applications of CRISPR technologies in research and beyond,” Nature biotechnology, vol. 34, no. 9, pp. 933–941, 2016.

[3] D.-K. Ro, E. M. Paradise, M. Ouellet, K. J. Fisher, K. L. Newman, J. M. Ndungu, K. A. Ho, R. A. Eachus, T. S. Ham, J. Kirby, et al., “Production of the antimalarial drug precursor artemisinic acid in engineered yeast,” Nature, vol. 440, no. 7086, pp. 940–943, 2006.

[4] J. Zhang, L. G. Hansen, O. Gudich, K. Viehrig, L. M. Lassen, L. Schrübbers, K. B. Adhikari, P. Rubaszka, E. Carrasquer-Alvarez, L. Chen, et al., “A microbial supply chain for production of the anti-cancer drug vinblastine,” Nature, vol. 609, no. 7926, pp. 341–347, 2022.

[5] B. Wang, R. I. Kitney, N. Joly, and M. Buck, “Engineering modular and orthogonal genetic logic gates for robust digital-like synthetic biology,” Nature communications, vol. 2, no. 1, p. 508, 2011.

[6] V. Hsiao, Y. Hori, P. W. Rothemund, and R. M. Murray, “A population-based temporal logic gate for timing and recording chemical events,” Molecular systems biology, vol. 12, no. 5, p. 869, 2016.

[7] M. Hicks, T. T. Bachmann, and B. Wang, “Synthetic biology enables programmable cell-based biosensors,” ChemPhysChem, vol. 21, no. 2, pp. 132–144, 2020.

[8] K. Wetterstrand, “DNA sequencing costs: Data from the NHGRI genome sequencing program (gsp) available at https://www.genome.gov/sequencingcostsdata,” Accessed August, 2019.

[9] R. A. Hughes and A. D. Ellington, “Synthetic DNA synthesis and assembly: putting the synthetic in synthetic biology,” Cold Spring Harbor Perspectives in Biology, vol. 9, no. 1, p. a023812, 2017.

[10] E. W. Sayers, M. Cavanaugh, K. Clark, K. D. Pruitt, C. L. Schoch, S. T. Sherry, and I. Karsch-Mizrachi, “Genbank,” Nucleic Acids Research, vol. 50, no. D1, p. D161, 2022.

[11] J. Jumper, R. Evans, A. Pritzel, T. Green, M. Figurnov, O. Ronneberger, K. Tunyasuvunakool, R. Bates, A. Žídek, A. Potapenko, et al., “Highly accurate protein structure prediction with alphafold,” Nature, vol. 596, no. 7873, pp. 583–589, 2021.

[12] A. A. Nielsen, B. S. Der, J. Shin, P. Vaidyanathan, V. Paralanov, E. A. Strychalski, D. Ross, D. Densmore, and C. A. Voigt, “Genetic circuit design automation,” Science, vol. 352, no. 6281, p. aac7341, 2016.

[13] A. J. Meyer, T. H. Segall-Shapiro, E. Glassey, J. Zhang, and C. A. Voigt, “Escherichia coli “marionette” strains with 12 highly optimized small-molecule sensors,” Nature Chemical Biology, vol. 15, no. 2, pp. 196–204, 2019.

[14] B. Canton, A. Labno, and D. Endy, “Refinement and standardization of synthetic biological parts and devices,” Nature Biotechnology, vol. 26, no. 7, pp. 787–793, 2008.

[15] C. Knief, A. Lipski, and P. F. Dunfield, “Diversity and activity of methanotrophic bacteria in different upland soils,” Applied and Environmental Microbiology, vol. 69, no. 11, pp. 6703–6714, 2003.

[16] J.-L. Ramos, M. Sol Cuenca, C. Molina-Santiago, A. Segura, E. Duque, M. R. Gómez-García, Z. Udaondo, and A. Roca, “Mechanisms of solvent resistance mediated by interplay of cellular factors in pseudomonas putida,” FEMS Microbiology Reviews, vol. 39, no. 4, pp. 555–566, 2015.

[17] H. E. Knights, B. Jorrin, T. L. Haskett, and P. S. Poole, “Deciphering bacterial mechanisms of root colonization,” Environmental Microbiology Reports, vol. 13, no. 4, pp. 428–444, 2021.

[18] T. Bjarnsholt, P. Ø. Jensen, M. J. Fiandaca, J. Pedersen, C. R. Hansen, C. B. Andersen, T. Pressler, M. Givskov, and N. Høiby, “Pseudomonas aeruginosa biofilms in the respiratory tract of cystic fibrosis patients,” Pediatric Pulmonology, vol. 44, no. 6, pp. 547–558, 2009.

[19] S. V. Nyholm and M. J. McFall-Ngai, “A lasting symbiosis: how the hawaiian bobtail squid finds and keeps its bioluminescent bacterial partner,” Nature Reviews Microbiology, vol. 19, no. 10, pp. 666–679, 2021.

[20] B. W. Biggs, S. R. Bedore, E. Arvay, S. Huang, H. Subramanian, E. A. McIntyre, C. V. Duscent-Maitland, E. L. Neidle, and K. E. J. Tyo, “Development of a genetic toolset for the highly engineerable and metabolically versatile Acinetobacter baylyi ADP1,” Nucleic Acids Research, vol. 48, no. 9, pp. 5169–5182, 2020.

[21] M. Taketani, J. Zhang, S. Zhang, A. J. Triassi, Y.-J. Huang, L. G. Griffith, and C. A. Voigt, “Genetic circuit design automation for the gut resident species bacteroides thetaiotaomicron,” Nature Biotechnology, vol. 38, no. 8, pp. 962–969, 2020.

[22] W. R. Whitaker, E. S. Shepherd, and J. L. Sonnenburg, “Tunable expression tools enable single-cell strain distinction in the gut microbiome,” Cell, vol. 169, no. 3, pp. 538–546, 2017.

[23] L. E. Leiva and A. Katz, “Regulation of leaderless mRNA translation in bacteria,” Microorganisms, vol. 10, Mar. 2022.

[24] S. D. Cole, A. E. Miklos, A. C. Chiao, Z. Z. Sun, and M. W. Lux, “Methodologies for preparation of prokaryotic extracts for cell-free expression systems,” Synthetic and Systems Biotechnology, vol. 5, no. 4, pp. 252–267, 2020.

[25] S. J. Moore, J. T. MacDonald, S. Wienecke, A. Ishwarbhai, A. Tsipa, R. Aw, N. Kylilis, D. J. Bell, D. W. McClymont, K. Jensen, et al., “Rapid acquisition and model-based analysis of cell-free transcription–translation reactions from nonmodel bacteria,” Proceedings of the National Academy of Sciences, vol. 115, no. 19, pp. E4340–E4349, 2018.

[26] S. J. Moore, H.-E. Lai, J. Li, and P. S. Freemont, “Streptomyces cell-free systems for natural product discovery and engineering,” Natural product reports, 2023.

[27] L. S. Thomashow and D. M. Weller, “Role of a phenazine antibiotic from pseudomonas fluorescens in biological control of gaeumannomyces graminis var. tritici,” Journal of bacteriology, vol. 170, no. 8, pp. 3499–3508, 1988.

[28] D. Haas and G. Défago, “Biological control of soil-borne pathogens by fluorescent pseudomonads,” Nature Reviews Microbiology, vol. 3, no. 4, pp. 307–319, 2005.

[29] D. L. McRose and D. K. Newman, “Redox-active antibiotics enhance phosphorus bioavailability,” Science, vol. 371, no. 6533, pp. 1033–1037, 2021.

[30] H. B. Klinke, A. Thomsen, and B. K. Ahring, “Inhibition of ethanol-producing yeast and bacteria by degradation products produced during pre-treatment of biomass,” Applied Microbiology and Biotechnology, vol. 66, pp. 10–26, 2004.

[31] J. Hirose, A. Nagayoshi, N. Yamanaka, Y. Araki, and H. Yokoi, “Isolation and characterization of bacteria capable of metabolizing lignin-derived low molecular weight compounds,” Biotechnology and bioprocess engineering, vol. 18, pp. 736–741, 2013.

[32] T. Kuhnigk and H. König, “Degradation of dimeric lignin model compounds by aerobic bacteria isolated from the hindgut of xylophagous termites,” Journal of Basic Microbiology, vol. 37, no. 3, pp. 205–211, 1997.

[33] I. Mahdi, N. Fahsi, M. Hijri, and M. Sobeh, “Antibiotic resistance in plant growth promoting bacteria: A comprehensive review and future perspectives to mitigate potential gene invasion risks,” Frontiers in Microbiology, vol. 13, 2022.

[34] R. Datta, A. Kelkar, D. Baraniya, A. Molaei, A. Moulick, R. S. Meena, and P. Formanek, “Enzymatic degradation of lignin in soil: a review,” Sustainability, vol. 9, no. 7, p. 1163, 2017.

[35] M. Bilal, S. Guo, H. M. N. Iqbal, H. Hu, W. Wang, and X. Zhang, “Engineering pseudomonas for phenazine biosynthesis, regulation, and biotechnological applications: a review,” World J. Microbiol. Biotechnol., vol. 33, p. 191, Oct. 2017.

[36] D. Haas and G. Défago, “Biological control of soil-borne pathogens by fluorescent pseudomonads,” Nat. Rev. Microbiol., vol. 3, pp. 307–319, Apr. 2005.

[37] K. Nesemann, S. A. Braus-Stromeyer, A. Thuermer, R. Daniel, D. V. Mavrodi, L. S. Thomashow, D. M. Weller, and G. H. Braus, “Draft genome sequence of the phenazine-producing Pseudomonas fluorescens strain 2-79,” Genome Announc., vol. 3, Mar. 2015.

[38] T. Timms-Wilson, R. Ellis, A. Renwick, D. Rhodes, D. Mavrodi, D. M. Weller, L. S. Thomashow, and M. Bailey, “Chromosomal insertion of phenazine-1-carboxylic acid biosynthetic pathway enhances efficacy of damping-off disease control by pseudomonas fluorescens,” Molecular Plant-Microbe Interactions, vol. 13, no. 12, pp. 1293–1300, 2000.

[39] J. P. Meier-Kolthoff and M. Göker, “TYGS is an automated high-throughput platform for state-of-the-art genome-based taxonomy,” Nature Communications, vol. 10, no. 1, p. 2182, 2019.

[40] P. I. Nikel, E. Martínez-García, and V. de Lorenzo, “Biotechnological domestication of pseudomonads using synthetic biology,” Nat. Rev. Microbiol., vol. 12, pp. 368–379, May 2014.

[41] Z. Z. Sun, C. A. Hayes, J. Shin, F. Caschera, R. M. Murray, and V. Noireaux, “Protocols for implementing an escherichia coli based TX-TL cell-free expression system for synthetic biology,” J. Vis. Exp., p. e50762, Sept. 2013.

[42] Z. Z. Sun, E. Yeung, C. A. Hayes, V. Noireaux, and R. M. Murray, “Linear DNA for rapid prototyping of synthetic biological circuits in an Escherichia coli based tx-tl cell-free system,” ACS Synthetic Biology, vol. 3, no. 6, pp. 387–397, 2014.

[43] M. Lanzer and H. Bujard, “Promoters largely determine the efficiency of repressor action.,” Proceedings of the National Academy of Sciences, vol. 85, no. 23, pp. 8973–8977, 1988.

[44] J. Meisner and J. B. Goldberg, “The Escherichia coli rhaSR-PrhaBAD inducible promoter system allows tightly controlled gene expression over a wide range in Pseudomonas aeruginosa,” Applied and environmental microbiology, vol. 82, no. 22, pp. 6715–6727, 2016.

[45] S. Zobel, I. Benedetti, L. Eisenbach, V. de Lorenzo, N. Wierckx, and L. M. Blank, “Tn7-based device for calibrated heterologous gene expression in Pseudomonas putida,” ACS Synthetic Biology, vol. 4, no. 12, pp. 1341–1351, 2015.

[46] K.-H. Choi and H. P. Schweizer, “mini-tn 7 insertion in bacteria with single att tn 7 sites: example Pseudomonas aeruginosa,” Nature Protocols, vol. 1, no. 1, pp. 153–161, 2006.

[47] K.-H. Choi, J. B. Gaynor, K. G. White, C. Lopez, C. M. Bosio, R. R. Karkhoff-Schweizer, and H. P. Schweizer, “A tn 7-based broad-range bacterial cloning and expression system,” Nature Methods, vol. 2, no. 6, pp. 443–448, 2005.

