## Supplemental Information for "Development of cell-free transcription-translation systems in three soil Pseudomonads"

June 9, 2023

**F. Time course data for *in vitro* TX-TL promoter panel reactions** **15**

**G. Time course data for *in vivo* promoter panel measurements** **16**

**H. Plasmid sequence** **17**

i. *P6d-mNeongreen* . . . . . . . . . . . . . . . . . . . . . . . . . 17

**I. Illustration of lysate production process** **18**

### A. Phylogenetic relationships and genome-to-genome distances

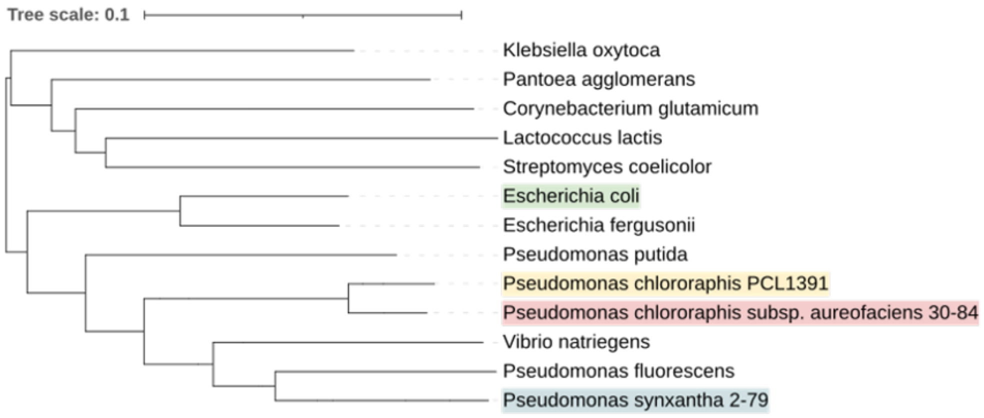

Supplementary Figure 1: Phylogenetic tree of bacteria used for in vitro transcription-translation (TX-TL), including the three in this study (highlighted). Figure was generated using the online iTOL tool [1].

| dDDH d4 in % [C.I.] (G+C difference) | *Pseudomonas synxantha* 2-79 | *Pseudomonas chlororaphis* PCL1391 | *Pseudomonas aureofaciens* 30-84 | *Escherichia coli* |
|---|---|---|---|---|
| *Pseudomonas synxantha* 2-79 | | 25.4 [23.0 - 27.8] (2.9%) | 25.6 [23.3 - 28.1] (3.17%) | 22.8 [20.5 - 25.2] (9.14%) |
| *Pseudomonas chlororaphis* PCL1391 | 25.4 [23.0 - 27.8] (2.9%) | | 58.3 [55.5 - 61.0] (0.27%) | 21.7 [19.5 - 24.1] (12.18%) |
| *Pseudomonas aureofaciens* 30-84 | 25.6 [23.3 - 28.1] (3.17%) | 58.3 [55.5 - 61.0] (0.27%) | | 21.8 [19.6 - 24.3] (12.31%) |
| *Escherichia coli* | 22.8 [20.5 - 25.2] (9.14%) | 21.7 [19.5 - 24.1] (12.18%) | 21.8 [19.6 - 24.3] (12.31%) | |

Supplementary Figure 2: Genome-to-genome distances as measured by digital DNA-DNA hybridization (dDDH), confidence intervals in [low - high] brackets and (G+C content difference) in parenthesis. The d4 measure is a genome-length independent metric described in [2]

### B. Growth rates of Pseudomonads

#### i. *P. synxantha*

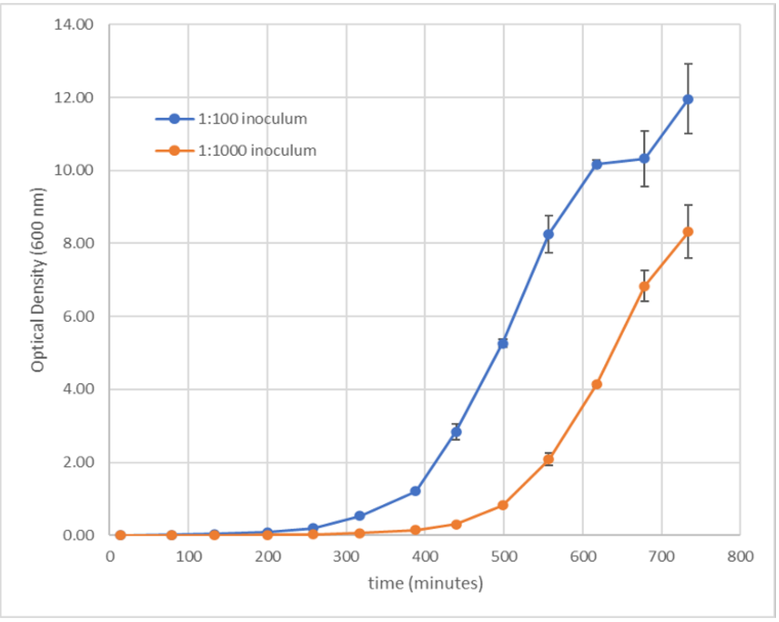

Supplementary Figure 3: Growth of *P. synxantha* 2-79 at 50 mL scale in 2xYTPG over 12 hours. For extract production, these cells were harvested at OD 3.0 +/- 10%. Error bars are standard deviation across cultures grown in triplicate.

#### 30 ii. *P. chlororaphis*

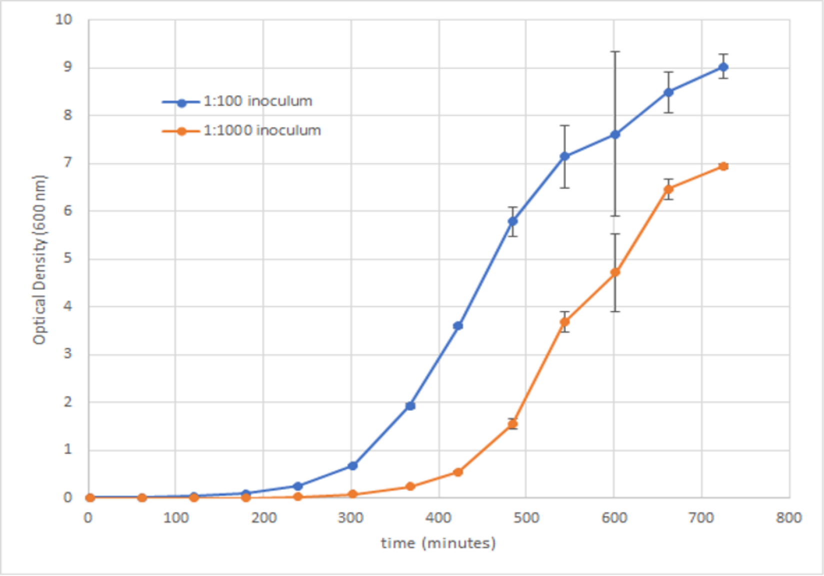

Supplementary Figure 4: Growth of *P. chlororaphis* PCL1391 at 50 mL scale in 2xYTPG over 12 hours. For extract production, these cells were harvested at OD 1.5 +/- 10%. Error bars are standard deviation across cultures grown in triplicate.

#### iii. *P. aureofaciens*

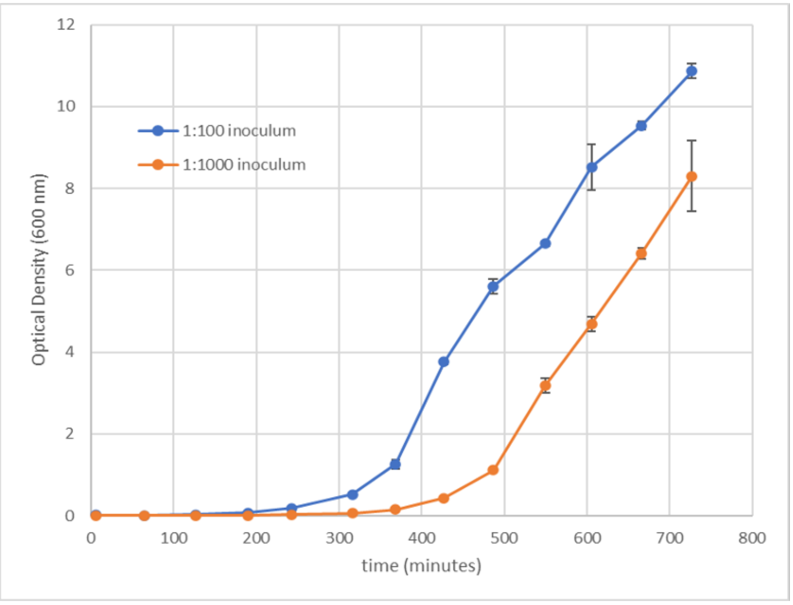

Supplementary Figure 5: Growth of *P. aureofaciens* 30-84 at 50 mL scale in 2xYTPG over 12 hours. For extract production, these cells were harvested at OD 3.0 +/- 10%. Error bars are standard deviation across cultures grown in triplicate.

#### C. Growth inhibitors in *E. coli* and *Pseudomonads*

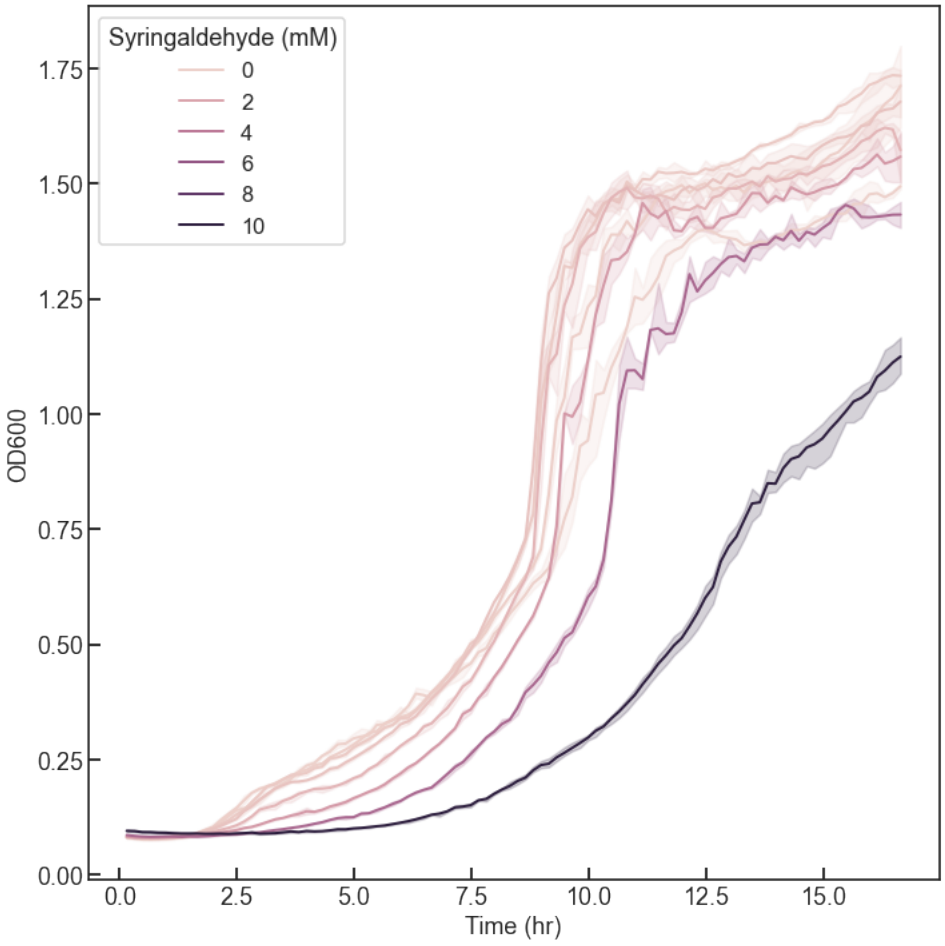

Supplementary Figure 6: A representative time course for *P. chlororaphis* growth in different concentrations of syringaldehyde.

### D. Protein concentrations of clarified lysates

#### i. *P. synxantha*

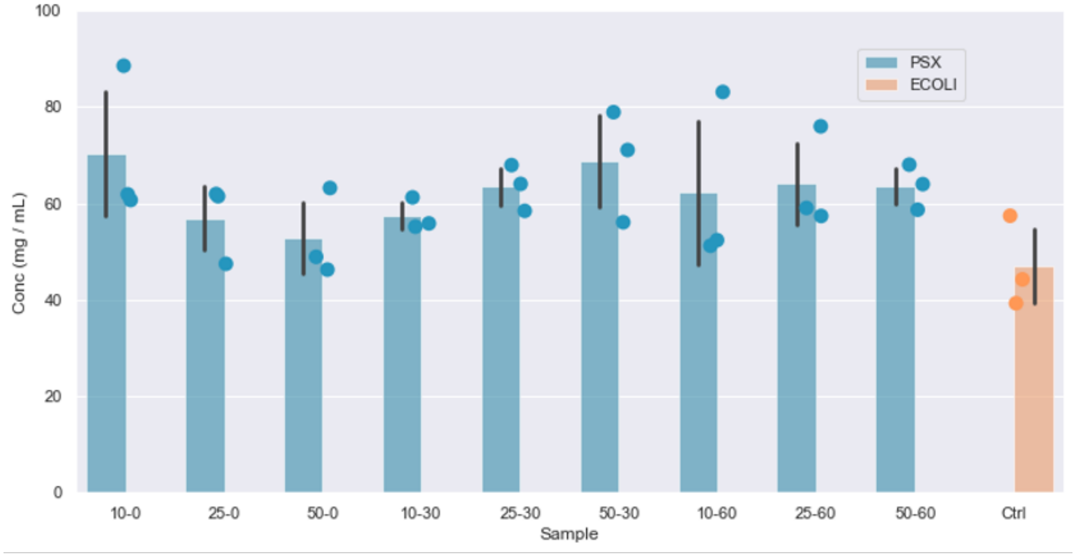

Supplementary Figure 7: Protein concentrations of different *P. synxantha* lysates as determined by Bradford assay. The sample labels at the bottom indicate “sonication amplitude” (left, arbitrary units) and “runoff time” (right, minutes). Bar heights are averages, error bars are standard deviation, and the dots are the individual measurements.

#### 36 ii. *P. chlororaphis*

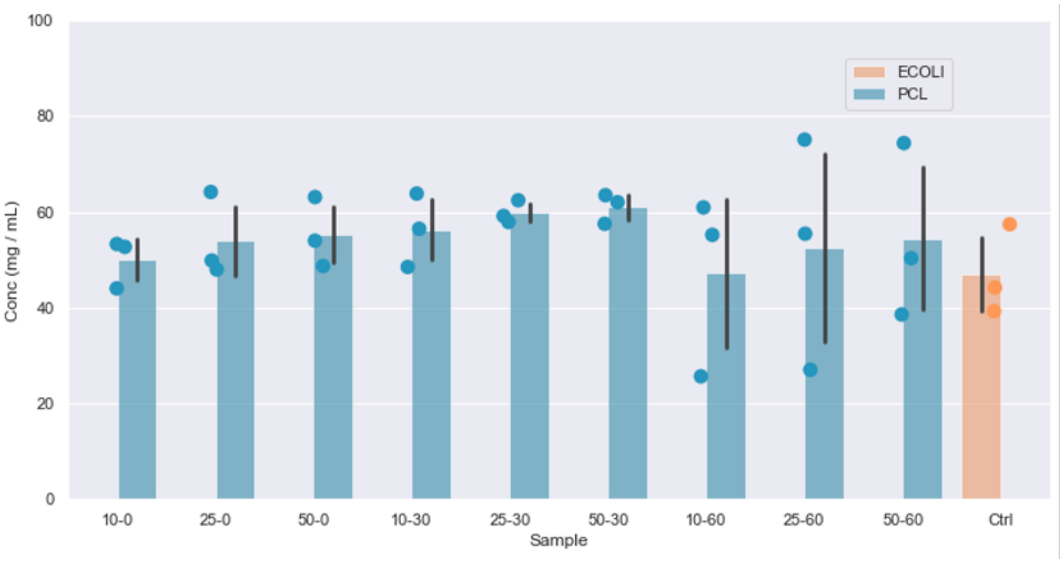

Supplementary Figure 8: Protein concentrations of different *P. chlororaphis* lysates as determined by Bradford assay. The sample labels at the bottom indicate "sonication amplitude" (left, arbitrary units) and "runoff time" (right, minutes). Bar heights are averages, error bars are standard deviation, and the dots are the individual measurements.

#### iii. *P. aureofaciens*

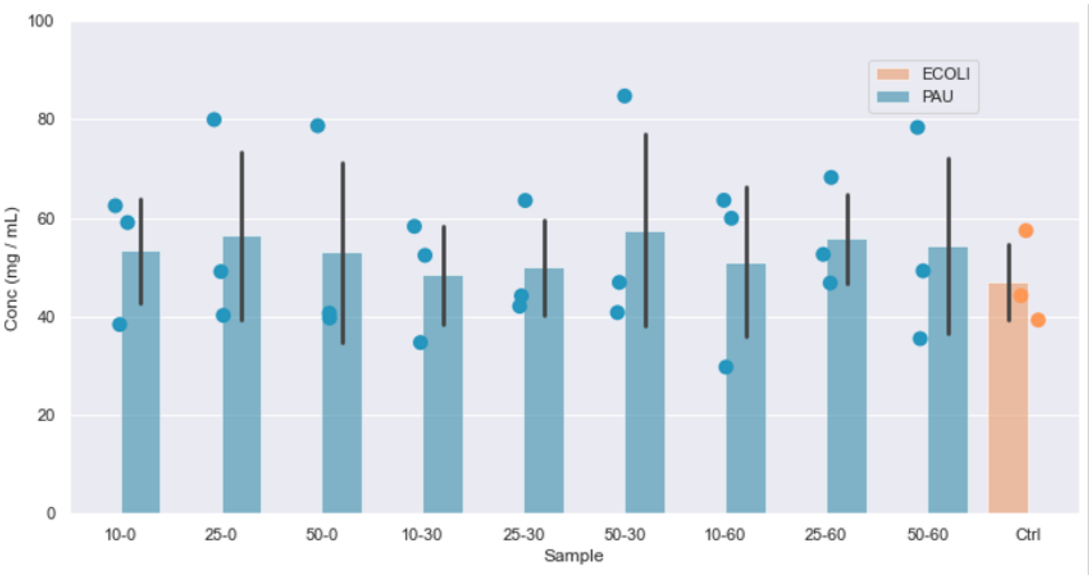

Supplementary Figure 9: Protein concentrations of different *P. aureofaciens* lysates as determined by Bradford assay. The sample labels at the bottom indicate “sonication amplitude” (left, arbitrary units) and “runoff time” (right, minutes). Bar heights are averages, error bars are standard deviation, and the dots are the individual measurements.

### E. Negative controls for selected reaction conditions

#### i. *P. synxantha*

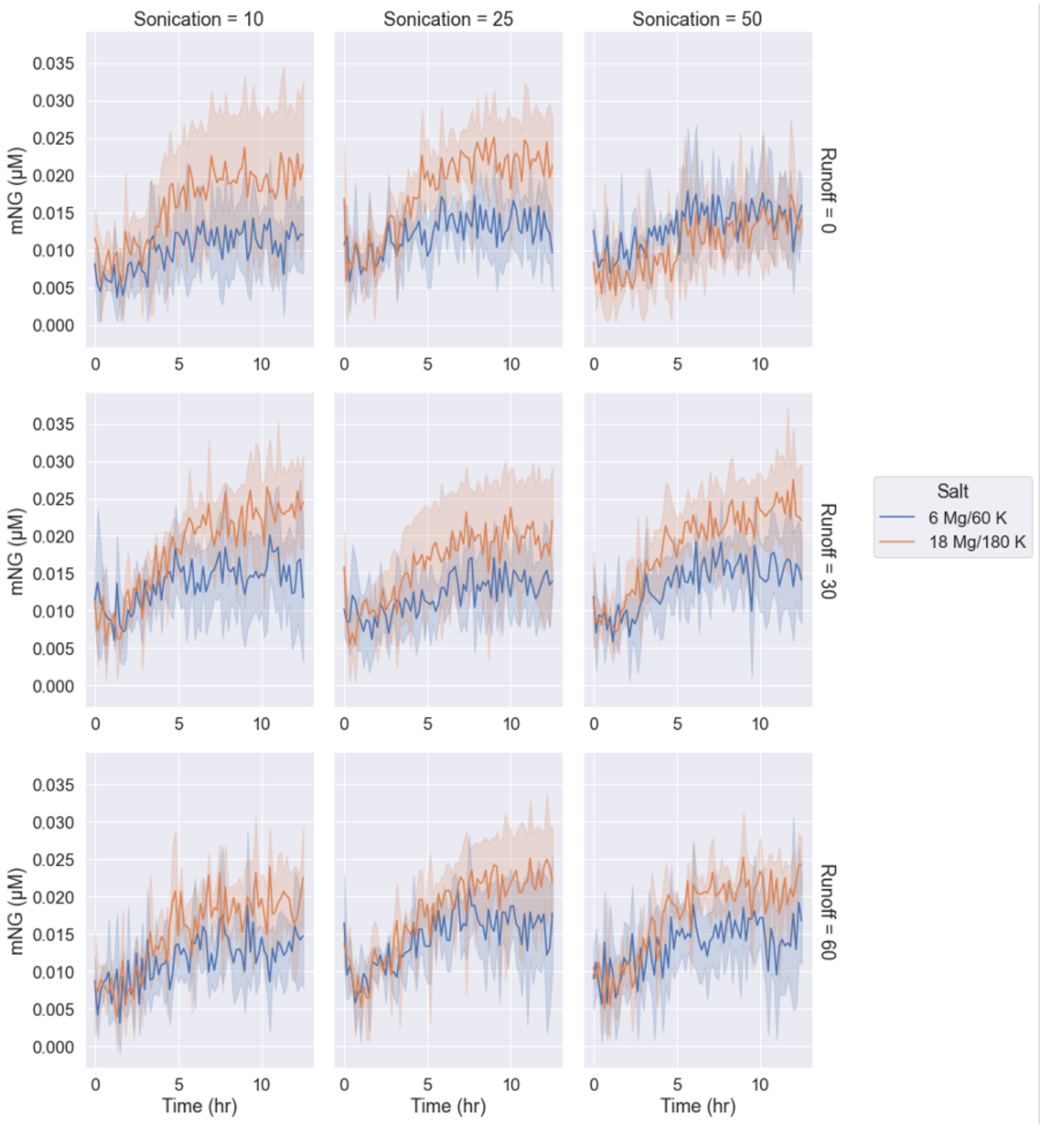

Supplementary Figure 10: Negative controls with no DNA template added to *P. synxantha* TX-TL reactions at two different salt concentrations, showing relatively low fluorescence and small changes in fluorescence over time in the mNeonGreen band (Ex: 490 nm, Em: 520 nm). The sample labels indicate “sonication amplitude” (top, arbitrary units) and “runoff time” (right, minutes).

#### ii. *P. chlororaphis*

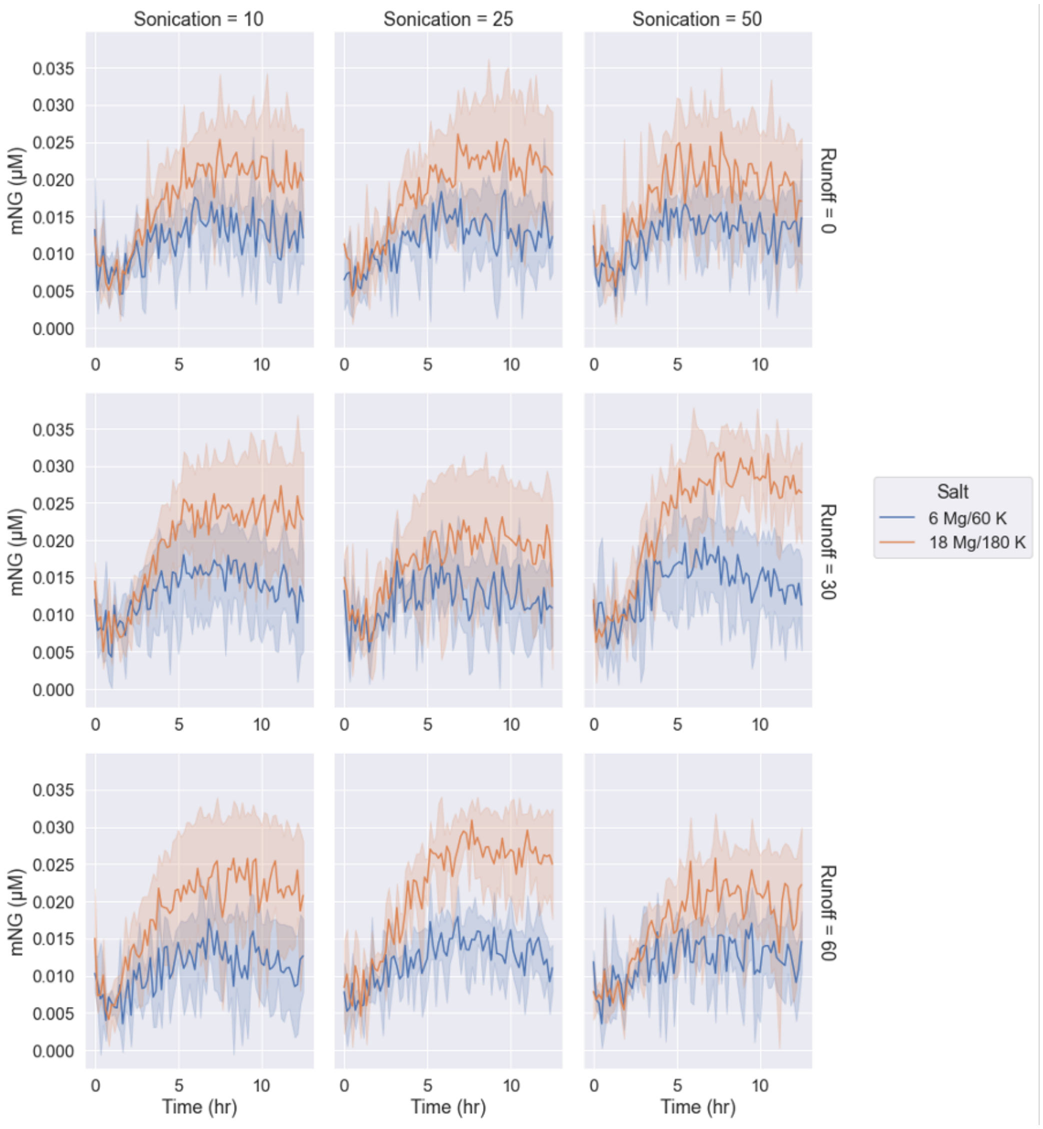

Supplementary Figure 11: Negative controls with no DNA template added to *P. chlororaphis* TX-TL reactions at two different salt concentrations, showing relatively low fluorescence and small changes in fluorescence over time in the mNeonGreen band (Ex: 490 nm, Em: 520 nm). The sample labels indicate "sonication amplitude" (top, arbitrary units) and "runoff time" (right, minutes).

42 ## iii. *P. aureofaciens*

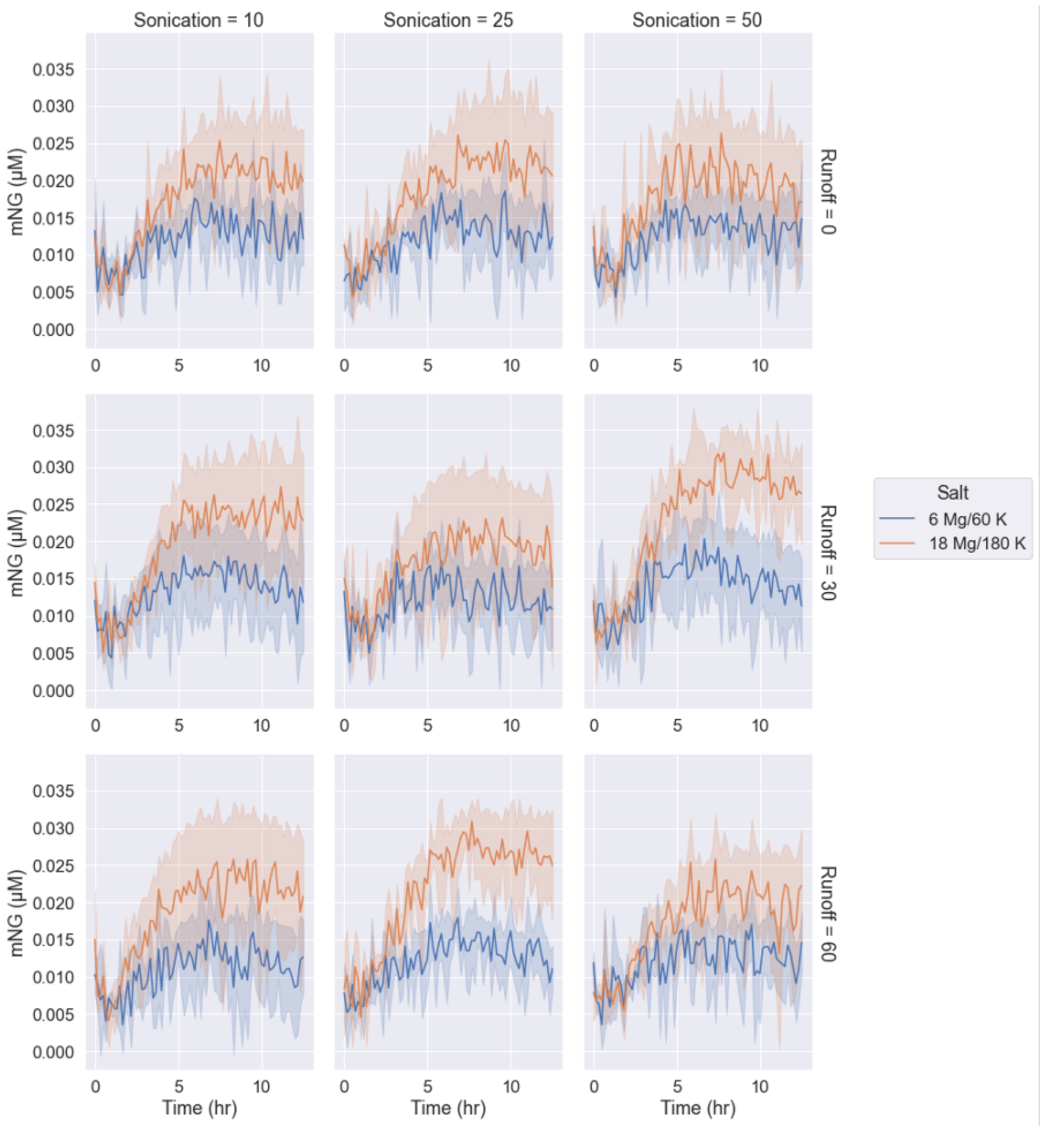

Supplementary Figure 12: Negative controls with no DNA template added to *P. aureofaciens* TX-TL reactions at two different salt concentrations, showing relatively low fluorescence and small changes in fluorescence over time in the mNeonGreen band (Ex: 490 nm, Em: 520 nm). The sample labels indicate “sonication amplitude” (top, arbitrary units) and “runoff time” (right, minutes).

#### F. Time course data for *in vitro* TX-TL promoter panel reactions

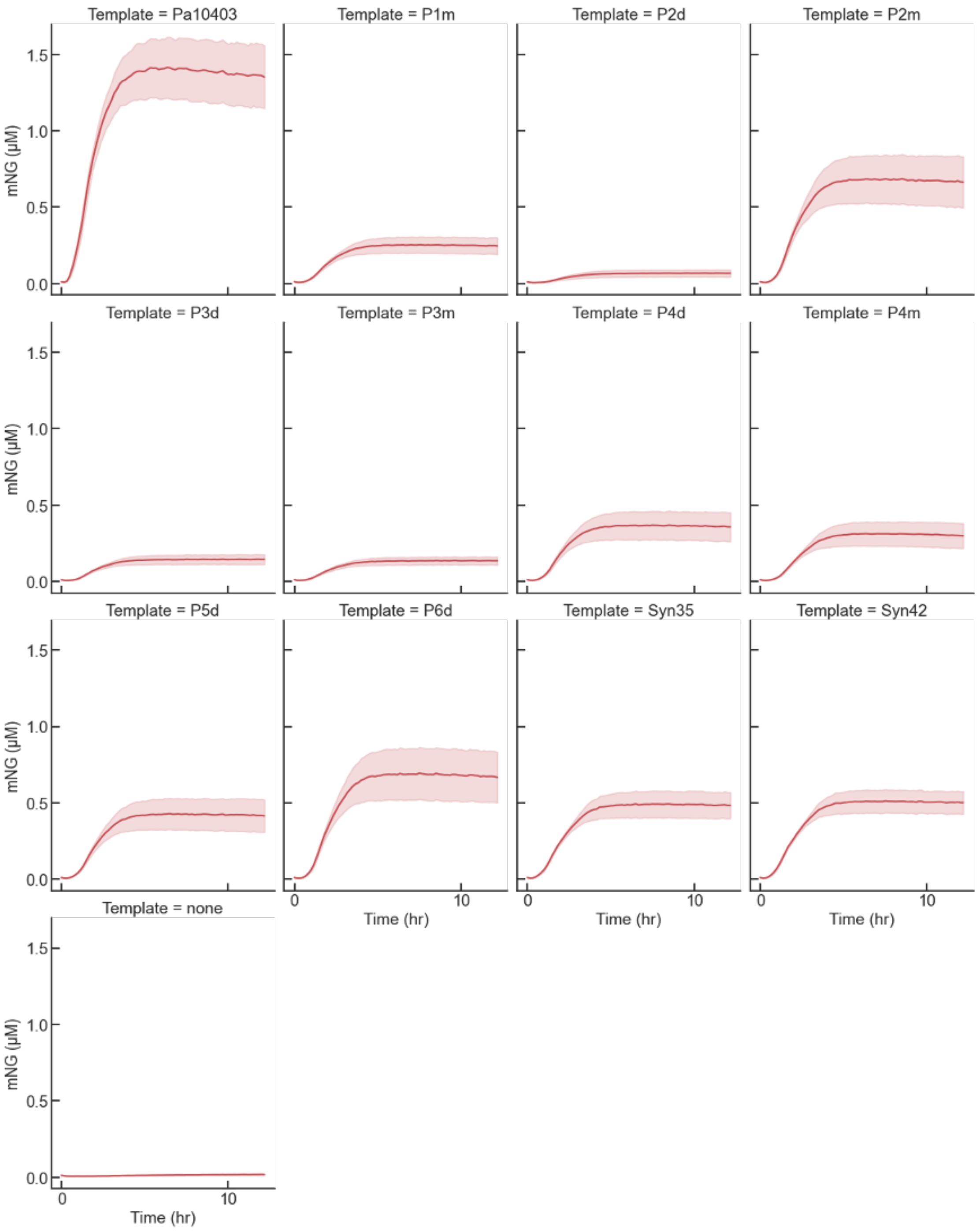

Supplementary Figure 13: Constitutive promoters driving the expression of mNeon-green during *P. synxantha* TX-TL reactions. The line shows the average of three biological replicates and the band shows the standard deviation.

### G. Time course data for *in vivo* promoter panel measurements

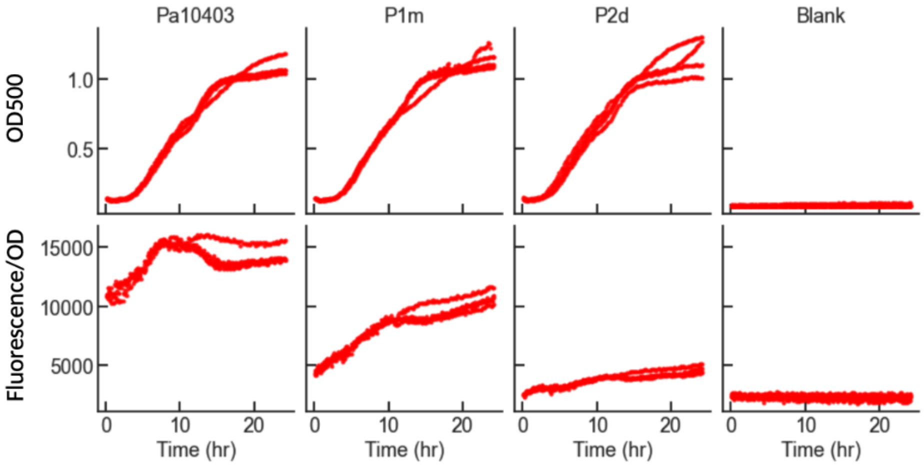

Supplementary Figure 14: Representative time courses showing genomically integrated constitutive promoters driving the expression of mNeongreen in living *P. synxantha* cells. All expression levels are normalized to the expression level of the strongest promoter fusion using the Pa10403 promoter. Each time course is showing one of three biological replicates and four technical replicates.

### H. Plasmid sequence

#### i. *P6d-mNeongreen*

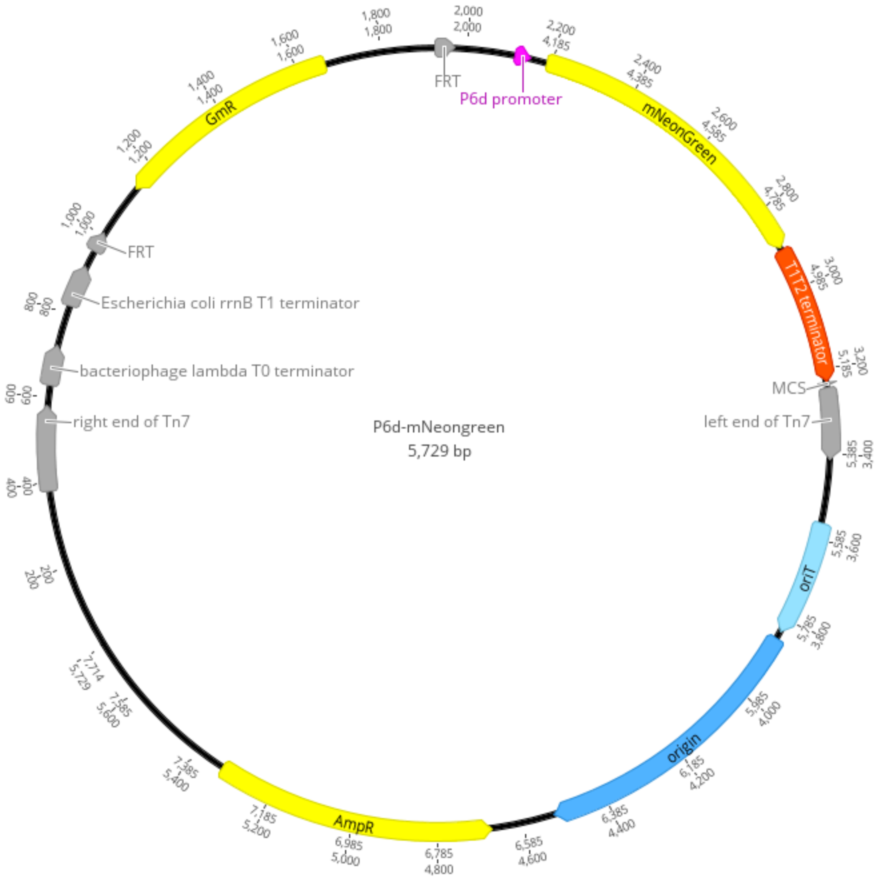

Supplementary Figure 15: Plasmid map of the pJM220 backbone with promoter P6d fused to mNeongreen. The map was generated using Geneious Prime v 2023.0

### I. Illustration of lysate production process

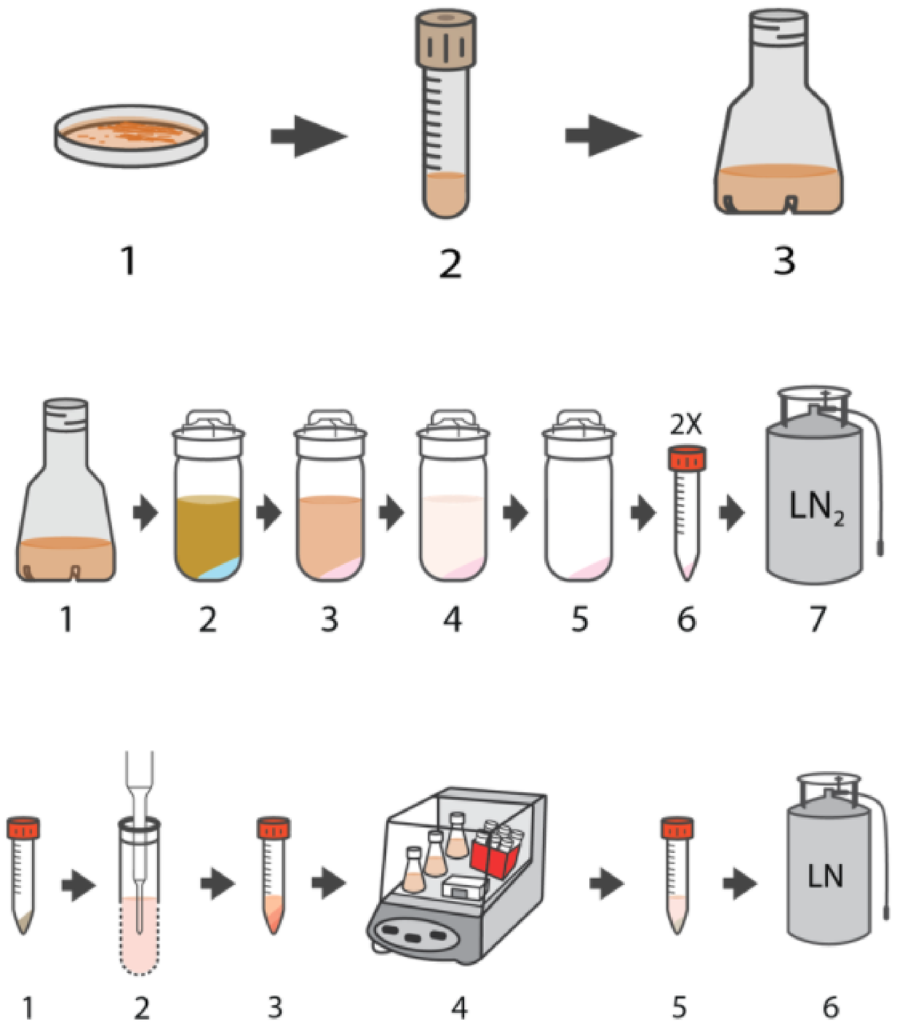

Supplementary Figure 16: Graphical illustration of the growth, harvest, extract production, and reaction set-up steps used in this study.

Briefly, in the top row step 1, cells are grown on 2xYTPG agar. After 18-36 hours of growth at 30C, the plates are stored at 4C for up to a month. In step 2, individual colonies are picked and used to grow 5 mL overnight cultures in 14 mL tubes. In step 3, 2.8L baffled shake flasks with 660 mL of 2xYTPG are inoculated 1:1000 the next day and grown for 8-12 hours. Note that unlike past protocols, the final culture is inoculated directly from a 5 mL overnight, without an intermediate 50 mL culture.

In the middle row step 1, cells are grown to OD 1.5-3.0, with monitoring of optical density performed every 60 minutes until the last doubling of the growth, then monitoring increases to once per 20 minutes. Dilutions of 1:4, 1:16, 1:64, or 1:256 in fresh 2xYTPG media are used to produce a sample with an optical density at 600 nm under 0.5. In step 2 200 mL of Buffer A (1.8 g/L Tris-acetate, 3 g/L Mg-acetate, and 12.2 g/L K-glutamate, adjusted to pH 8.2 with 2M Tris and autoclaved, then 1 mL / L of 1M DTT is added) is frozen in 1L centrifuge tubes stored at -20C at an angle for 1-2 hours.

At harvest time, the cell culture is decanted into the 1L centrifuge tube over the ice. This brings the culture temperature down from 30C to 10C within a few minutes. In step 3, the cell culture is centrifuged at 4800g for 12 minutes at 4C. The supernatant is decanted, and the tubes are placed on ice. The cell pellet is resuspended in Buffer A, then otherwise follows the protocol in [3] with two wash steps in 1L bottles, followed by transfer into two weighed 50 mL tubes and a final wash step. The supernatant is removed and the pellet is flash frozen in liquid nitrogen and stored at -80C.

At a later time, the cell pellet is removed from the freezer and placed on wet ice to thaw. In the bottom row step 1, 1 mL / g of Buffer A is used to resuspend the pellet, then 4 mL of resuspended cell pellet is added to each 14 mL Falcon tube. In step 2, each tube is sonicated for 120s, 5s on, 10s off, with an amplitude of 10, 25, or 50 on a Qsonica Q700. In step 3, the tubes are centrifuged at 12,000g for 10 minutes at 4C. In step 4, the supernatant is transferred to new 1.4 mL or 14 mL tubes and incubated in a “runoff” step with open lids shaking at 220 rpm at 30C for 0-60 minutes. These tubes are centrifuged again at 12,000g for 10 minutes at 4C, then the supernatant is transferred to a new tube. Aliquots of this final clarified lysate are made for freezing in LN2 and storage at -80C.

### References

[1] I. Letunic and P. Bork, "Interactive tree of life (itol) v5: an online tool for phylogenetic tree display and annotation," *Nucleic acids research*, vol. 49, no. W1, pp. W293–W296, 2021.

[2] J. P. Meier-Kolthoff and M. Göker, "TYGS is an automated high-throughput platform for state-of-the-art genome-based taxonomy," *Nat. Commun.*, vol. 10, p. 2182, May 2019.

[3] Z. Z. Sun, C. A. Hayes, J. Shin, F. Caschera, R. M. Murray, and V. Noireaux, "Protocols for implementing an escherichia coli based TX-TL cell-free expression system for synthetic biology," *J. Vis. Exp.*, p. e50762, Sept. 2013.
